# Occlusion of lip movements impairs reconstruction of acoustic speech features and higher-level segmentational features in the presence of a distractor speaker

**DOI:** 10.1101/2021.09.28.461909

**Authors:** Chandra Leon Haider, Nina Suess, Anne Hauswald, Hyojin Park, Nathan Weisz

**Affiliations:** Centre for Cognitive Neuroscience and Department of Psychology, University of Salzburg, Austria; School of Psychology & Centre for Human Brain Health (CHBH), University of Birmingham, Birmingham, UK; Neuroscience Institute, Christian Doppler University Hospital, Paracelsus Medical University, Salzburg, Austria

**Keywords:** stimulus reconstruction, face masks, audiovisual speech, formants, speech envelope

## Abstract

Multisensory integration enables stimulus representation even when the sensory input in a single modality is weak. In the context of speech, when confronted with a degraded acoustic signal, congruent visual inputs promote comprehension. When this input is occluded speech comprehension consequently becomes more difficult. But it still remains inconclusive which levels of speech processing are affected under which circumstances by occlusion of the mouth area. To answer this question, we conducted an audiovisual (AV) multi-speaker experiment using naturalistic speech. In half of the trials, the target speaker wore a (surgical) face mask, while we measured the brain activity of normal hearing participants via magnetoencephalography (MEG). We additionally added a distractor speaker in half of the trials in order to create an ecologic difficult listening situation. A decoding model on the clear AV speech was trained and used to reconstruct crucial speech features in each condition. We found significant main effects of face masks on the reconstruction of acoustic features, such as the speech envelope and spectral speech features (i.e. pitch and formant frequencies), while reconstruction of higher level features of speech segmentation (phoneme and word onsets) were especially impaired through masks in difficult listening situations. As we used surgical face masks in our study, which only show mild effects on speech acoustics, we interpret our findings as the result of the occluded lip movements. This idea is in line with recent research showing that visual cortical regions track spectral modulations. Our findings extend previous behavioural results, by demonstrating the complex contextual effects of occluding relevant visual information on speech processing.

**Highlights:**

- Surgical face masks impair neural tracking of speech features
- Tracking of acoustic features is generally impaired, while higher level segmentational features show their effects especially in challenging listening situations
- An explanation is the prevention of a visuo-phonological transformation contributing to audiovisual multisensory integration

## Introduction

Despite being initially processed by different sensory organs and brain regions, information from different modalities are used to build a coherent perceptual experience. As sources of sensory inputs are usually multi-modal in natural environments, the brain has developed mechanisms to utilize information from one modality in order to facilitate processing of another. Focusing on vision and audition, past studies showed responses to regular visual patterns in the auditory cortex, indicating a modulatory influence of the early visual regions on primary auditory regions (Suess et al., 2021; for a review see: Bauer et al., 2020). This process is not restricted to simple stimuli such as tone sequences or light flashes, but can be observed in more complex stimuli like speech (Peelle & Sommers, 2015). After establishing audiovisual enhancement of speech comprehension behaviorally (Sumby & Pollack, 1954), more recent studies followed up on the research of simple stimuli and tried to explore audiovisual speech processing to the neural level (Crosse et al., 2015; Crosse, Di Liberto, & Lalor, 2016; Golumbic et al., 2013; Park et al., 2016, Peele & Sommer, 2015). One promising approach to investigate these effects is the use of decoding models. Hereby, the participant’s brain response obtained using electroencephalography (EEG) or magnetoencephalography (MEG) is linked to the presented stimulus in order to measure how well a certain feature is encoded in the brain (Crosse, Di Liberto, Bednar, et al., 2016). Using this and similar approaches, past studies showed that the brain directly tracks speech specific components like the speech envelope (Brodbeck & Simon, 2020; Ding & Simon, 2014). More importantly, a study demonstrated that in challenging listening situations, audiovisual gain can be quantified not only on the behavioral level, but can also be assessed on a neural level by using the tracking of the speech envelope as proxy for speech comprehension (Crosse, Di Liberto, & Lalor, 2016). These results are in line with the concept called “inverse effectiveness” (Meredith & Stein, 1986), stating that multisensory gain is increased when individual modalities are of low quality compared to an ideal environment (i.e. visual gain is increased when the acoustic speech signal is noisy in a audiovisual (AV) listening situation).

On the one hand, this mentioned audiovisual facilitation might be explained by providing simple temporal cues (i.e. opening and closing of mouth) when having to attend to auditory stimuli (Van Engen et al., 2019). On the other hand, visual information might be preselecting certain possible stimuli (e.g. phonemes) and therefore enhancing subsequent auditory processing as a form of crossmodal integration. By using the additive model (i.e. comparing event-related potentials (ERP) to audio stimuli + ERPs visual stimuli (A+V) to ERPs of audiovisual stimuli (AV)), past studies indeed suggested that the brain integrates early information from the visible lip movements in the auditory cortex for efficient speech processing (Besle et al., 2004, 2009).

In addition to these effects in auditory processing regions, we have provided evidence for a direct visuo-phonological transformation when individuals only process visual information (i.e. silent video recordings of speakers), by showing that the acoustic speech envelope is tracked in visual cortical regions when individuals observe lip movements (Hauswald et al., 2018; Suess, Hauswald, et al., 2021). Furthermore, when again provided with only visual speech, the visual cortex also tracks spectral modulations in the range of the pitch, as well as in the second (F2) and third formant (F3), which reflect mainly sounds produced with the visible part of the mouth (Suess, Hauswald, et al., 2021). These results align well with previous findings by Chandrasekaran et al. (2009), who indicate that the area of mouth opening correlates strongest with spectral components of speech in the range of 1 kHz - 3 kHz, corresponding to the frequency range of F2 and F3. Another study highlighting the importance of formants was published by Plass and colleagues (2020). They showed an even stronger audiovisual enhancement through formant frequencies than the well established speech envelope. Together, these results reveal that visual lip movements are transformed in order to track acoustic speech features such as the speech envelope and formant frequencies leading to efficient processing, especially when acoustics are distorted. These findings are even more important when put into the context of the ongoing Covid-19 pandemic, as the use of face masks as an effective intervention against aerosol transmission could at the same time hinder the integration of visual and acoustic information for optimal speech understanding.

Indeed, a large online study investigated effects of face masks on audiovisual (AV) speech comprehension behaviourally (Brown et al., 2021). They found no differences in sentence intelligibility between clear AV speech (i.e. no face mask) and face masks of several types (e.g. surgical face mask and N95 mask) in conditions with a quiet background, but differences became apparent in conditions with moderate and high background noise. Despite these well-established effects, the behavioural studies have left open which (degraded) speech features are driving these findings. Decoding distinct speech features from the neural signal could be used for addressing this issue. Putting the aforementioned findings together, face masks might adversely impact the ability to integrate visual and auditory information from diverse speech characteristics at different hierarchical levels, resulting in poor behavioural performance. With face masks still common in everyday life as a measure against Covid-19 and continuing to remain important in medical settings, understanding precisely which features of speech are less well tracked by the brain can help guide decisions on which face mask to use. These considerations are especially important when dealing with hearing-impaired individuals (Puschmann et al., 2019).

In the current MEG study, we investigated how neural tracking of a variety of speech features (purely acoustic and lexical/phonetic boundaries) in an audio-visual naturalistic speech paradigm is impaired through (surgical) face masks. Special emphasis is placed on an interaction between face masks and difficult listening situations induced via an audio-only distractor speaker, as studies emphasised the visual benefit when acoustics are unclear (Brown et al., 2021; Crosse, Di Liberto, & Lalor, 2016; Mitchel & Weiss, 2014; Park et al., 2016; Sumby & Pollack, 1954). We then trained a backward model on clear speech in order to reconstruct the speech characteristics from the participants’ brain data for each condition. Additionally, we measured participants’ comprehension performance and subjective difficulty ratings. We hypothesised strong effects on speech features that have been shown to be detectable by visual input (i.e. lip movements). Such features are the speech envelope, pitch, the averaged F2 and F3 (F2/3) as well as segmentational features (i.e. phoneme and word onsets), because here multisensory gain can be expected for natural AV speech. We found strong adverse general effects of the face mask on acoustic feature (i.e. speech envelope and spectral features) reconstruction and difficulty ratings irrespective of a distractor speaker. Importantly, for features of segmentation (word and phoneme onsets) face masks revealed their adverse impact especially in difficult listening situations.

## Methods

### Participants

29 German native speakers (12 female) aged between 22 and 41 years (*M = 26*.*79, SD = 4*.*86*) took part in our study. All participants had self-reported normal hearing, verified by a standard clinical audiometry. Further exclusion criteria were non-removable magnetic objects, as well as a history of psychiatric or neurological conditions. Recruitment was done via social media and university lectures. One participant was excluded because signal source separation could not be applied to the MEG dataset. All participants signed an informed consent form and were compensated with €10 per hour or course credit. The experimental protocol was approved by the ethics committee of the University of Salzburg and was carried out in accordance with the Declaration of Helsinki.

### Stimuli

We used excerpts of four different stories for our recording read out in German. ‘Die Schokoladenvilla - Zeit des Schicksals. Die Vorgeschichte zu Band 3’ (“The Chocolate Mansion, The Legacy” – prequel of Volume 3”) by Maria Nikolai and ‘Die Federn des Windes’ (“The feathers of the wind”) by Manuel Timm were read out by a female speaker. ‘Das Gestüt am See. Charlottes großer Traum’ (“The stud farm by the lake. Charlotte’s great dream”) by Paula Mattis and ‘Gegen den Willen der Väter’ (“Against the will of their fathers”) by Klaus Tiberius Schmidt were read out by a male speaker.

Stimuli were recorded using a Sony FS100 camera with a sampling rate of 25 Hz and a Rode NTG 2 microphone with a sampling rate of 48 kHz. We aimed at a duration for each story of approximately ten minutes, which were cut into ten videos of around one minute each (*range: 56 s - 76 s, M = 64 s, SD = 4*.*8 s)*. All stories were recorded twice, once without the speaker wearing a surgical face mask and once with the speaker wearing a surgical face mask (Type IIR, three-layer single-use medical face mask, see Figure 1A). After cutting the videos, we ended up with 80 videos of approximately one minute each. Forty of those were presented to each participant (20 with a female speaker, 20 with a male speaker) in order to rule out sex-specific effects. The audio track was extracted and stored separately. The audio files were then normalised using the Python function ‘ffmpeg-normalise’ with default options. Pre-recorded audiobooks read out by different speakers (one female, one male) were used for the distractor speaker and normalised using the same method. These audio files contained either a (different) single male or female speaker. The syllable rate was analysed using a Praat script (Boersma & Weenink, 2001; de Jong & Wempe, 2009). The target speakers’ syllable rates varied between 3.7 Hz and 4.6 Hz (*M = 4*.*1 Hz*). Target and distractor stimuli were all played to the participant at the same volume, which was individually set to a comfortable level at the start of the experiment.

**Figure 1.**
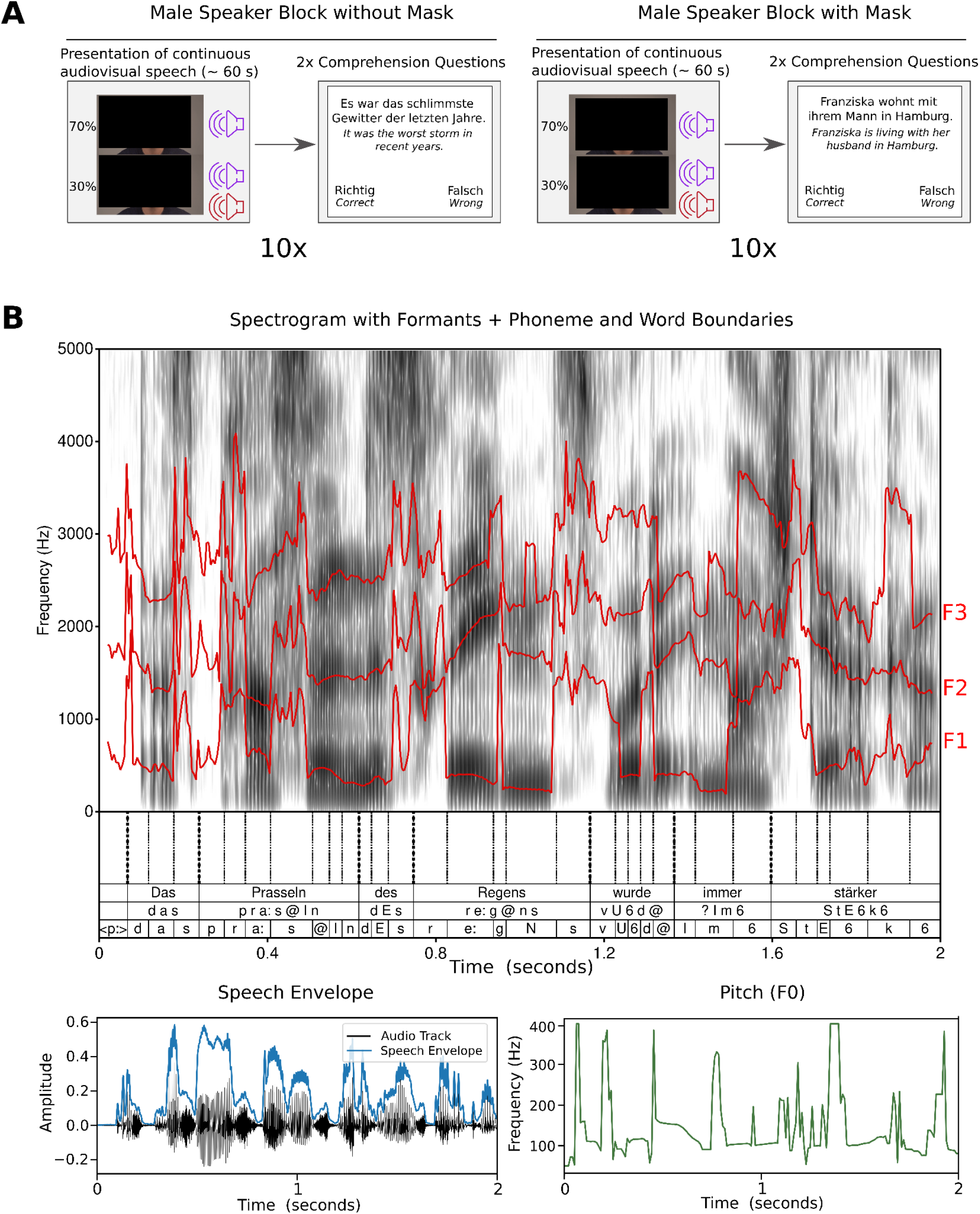
Experimental procedure and speech features. **A** shows two example blocks with a male target speaker. In the block on the left the speaker did not wear a face mask across the ten trials per block. In 70% of trials the target speaker was presented solitarily, in 30% a same sex audio-only distractor speaker was added at the same volume (denoted by the second sound icon). After each of the ten trials per block, two ‘true or false’ comprehension questions were presented to the participant (italic letters underneath depict English translation). Participants answered via button press (left or right button). On the right, a block is depicted with the male speaker wearing a face mask across the ten trials of the block. Otherwise the procedure is the same as the block without a face mask. Clear speech is defined as the condition without a mask and without a distractor speaker. The two depicted blocks were repeated with a female speaker, resulting in a total of four blocks. **B** shows the speech features investigated. Formants (F1 - F3) are shown in red overlaid on the speech spectrogram. Segmentation in phonemes and words (top row: orthographic word; mid row: phonetic word; bottom row: phoneme) was done using forced alignment. This segmentation can be seen on the bottom of the spectrogram. The speech envelope can be seen on the bottom left of the figure. On the bottom right of the figure, the speaker’s pitch or fundamental frequency (F0) is depicted. All depictions are based on the same two-second long speech interval.

<u>* images have been removed/obscured due to a bioRxiv policy on the inclusion of faces.</u>

### Experimental procedure

Before the start of the experiment, we performed a standard clinical audiometry using a AS608 Basic (Interacoustics, Middelfart, Denmark) in order to assess participants’ individual hearing ability. Afterwards, participants were prepared for MEG (see *Data acquisition*).

We started the MEG measurement with five minutes of resting-state activity (not included in this manuscript). We then assessed the participants’ individual hearing threshold in order to adjust our stimulation volume. If the participant stated afterwards that stimulation was not comfortable or not loud enough, we adjusted the volume again manually to the participant’s requirement. Of the four stories, half were randomly chosen with the target speakers wearing face masks in the recording. In the remaining half, speakers did not wear a face mask. Each story presentation functioned as one stimulation block, resulting in four blocks overall. One block consisted of ten ∼ 1 minute long trials. In three randomly selected trials per block-(i.e. 30% of trials), a same-sex audio-only distractor speaker was added at equal volume as the target speaker. We only added a distractor speaker in 30% of trials in order to retain enough data to train our backward model on clear speech (see stimulus reconstruction section). Distractor speaker presentation started five seconds after target speaker video and audio onset in order to give the participants time to pay attention to the target speaker. Within the blocks, the story presentation followed a consistent storyline across trials. After each trial, two unstandardised ‘true or false’ statements regarding semantic content were asked to assess comprehension performance and keep participants focused (Figure 1A). Additionally, participants rated subjective difficulty and motivation at four times per block on a five-point likert scale (not depicted in Figure 1A). The participants’ answers were given via button presses. In one half of the four blocks a female target speaker was presented, in the other half a male target speaker. Videos were back-projected on a translucent screen with a screen diagonal of 74 cm via a Propixx DLP projector (Vpixx technologies, Canada) ∼ 110 cm in front of the participants. It was projected with a refresh rate of 120 Hz and a resolution of 1920 × 1080 pixels. Including preparation, the experiment took about 2 hours per participant. The experiment was coded and conducted with the Psychtoolbox-3 (Brainard, 1997; Kleiner et al., 2007; Pelli, 1997) with an additional class-based library (‘Objective Psychophysics Toolbox’, o_ptb) on top of it (Hartmann & Weisz, 2020).

### Data acquisition

We recorded brain data with a sampling rate of 1 kHz at 306-channels (204 first-order planar gradiometers and 102 magnetometers) with a Triux MEG system (MEGIN, Helsinki, Finnland). The acquisition was performed in a magnetically shielded room (AK3B, Vacuumschmelze, Hanau, Germany). Online bandpass filtering was performed from 0.1 Hz to 330 Hz. Prior to the acquisition, cardinal head points (nasion and pre-auricular points) were digitised with a Polhemus FASTTRAK Digitizer (Polhemus, Colchester, Vermont, USA) along with around 300 points on the scalp in order to assess individual head shapes. Using a signal space separation algorithm provided by the MEG manufacturer (Maxfilter, version 2.2.15), we filtered noise resulting from sources outside the head and realigned the data to a standard head position, which was measured at the beginning of each block.

### Speech feature extraction

All the speech features investigated are depicted in Figure 1B.The speech envelope was extracted using the Chimera toolbox. For this purpose, the speech signal was filtered forward and in reverse with a 4th order Butterworth bandpass filter at nine different frequency bands equidistantly spaced between 100 and 10000 Hz corresponding to the cochlear map (Smith et al., 2002). Then, a Hilbert transformation was performed to extract the envelopes from the resulting signals. These nine envelopes were then summed up to one general speech envelope and normalised.

The pitch (fundamental frequency, F0) was extracted using the built-in Matlab Audio toolbox function *pitch*.*m* and downsampled to 50 Hz. The speech formants (first, second, third and the averaged second and third formant) were extracted using FormantPro (Xu & Gao, 2018), a tool for automatic formant detection via Praat (Boersma & Weenink, 2001) at 50 Hz with an integration window length of 20 ms, and a smoothing window of 10 ms length.

Phoneme and word onset values were generated using forced alignment with MAUS web services (Kisler et al., 2017; Schiel, 1999) in order to obtain a measure for speech segmentation. We generated two time-series with binary values indicating an onset of phoneme or word, respectively. Then, we smoothed the time-series of binary values using a gaussian window with a width of 10 ms. In the end, all features were downsampled to 50 Hz to match the sampling rate of the corresponding brain signal, as most speech relevant signals present themselves below 25 Hz (Crosse et al., 2021).

### MEG preprocessing

The raw data was analysed using Matlab R2020b (The MathWorks, Natick, Massachusetts, USA) and the FieldTrip toolbox (Oostenveld et al., 2011). First, we computed 50 independent components to remove eye and heart artifacts. We removed on average 2.38 components per participant (*SD =* .*68*). We further filtered the data using a sixth-order zero-phase Butterworth bandpass filter between 0.1 and 25 Hz. Afterwards, we epoched the data into 2.5 s segments. Finally, we downsampled our data to 50 Hz.

### Stimulus reconstruction

To reconstruct the different speech characteristics (speech envelope, pitch, resonant frequencies as well as word and phoneme onsets) from the brain data, we used the mTRF Toolbox (Crosse, Di Liberto, Bednar, et al., 2016). The goal of this approach is to map brain responses (i.e. all 306 MEG channels) back to the stimulus(-feature) (e.g. speech envelope) using linear models in order to obtain a measure of how well a certain characteristic is encoded in the brain. According to our 2×2 experimental design, the stimulus features were reconstructed for each condition. As the distractor speaker starts after five seconds of the trial start, these five seconds were not assigned to the *Distractor* condition, but rather reassigned to their respective condition with only a single speaker.

The stimulus features and the brain data at all 306 MEG channels were z-scored and the epochs were shuffled. We then used the clear speech condition (with no masks and no distractor speaker presented) to train the backward model with ridge regression. In order to test the model on a clear audio data set as well, we split it into seven parts and trained our model on six parts, while using the remaining part to test it. This results in approximately twelve minutes of data for training the model. We defined our time lags to train our model from -150 ms to 450 ms. Then, we performed seven-fold leave-one-out cross-validation on our training dataset to find the optimal regularisation parameter (Willmore & Smyth, 2003) in the range of 10^0^… 10^5^. We used the same data with the obtained regularisation parameter to train our backward model. For each condition, we used the same backward model trained on clear speech to reconstruct the speech characteristics of interest, namely the speech envelope, pitch, resonant frequencies (F1-3 and F2/3) and segmentational features (phoneme and word onsets). As we used clear audio trials for training the decoding model and added a distractor speaker only in 30% percent of trials (see *Experimental procedure*, Figure 1A), this resulted in a variable length of test data sets. In the ‘no mask/no distractor’ condition it was ∼ 2 minutes, in the ‘mask/no distractor’ condition it was ∼ 14 minutes and for ‘no mask/distractor’ as well as ‘mask/distractor’ condition it was ∼ 6 minutes each. The process was repeated six times, so that each subset of the clear speech condition was used as a test set while all other subsets were used for training. For each participant, each speech feature and each of the four conditions we computed the correlation coefficient (Pearson’s *r*) of the reconstructed feature and the original feature as a measure of reconstruction accuracy. This was done by Fisher z-transformation and averaging all respective correlation coefficients for each test set and each of the seven repetitions obtained through the aforementioned procedure.

### Statistical analysis

We performed a repeated measures ANOVA with the within-factors *Mask* (no face mask vs. face mask) and *Distractor* (no distractor speaker vs. distractor speaker) and the obtained Fisher z-transformed correlation coefficients (i.e. reconstruction accuracy) as dependent variables.

For the behavioural results (comprehension performance and subjective difficulty), we also used a repeated measures ANOVA with the same factors *Mask* and *Distractor*. We used comprehension performance scores (i.e. the percentage of correct answers) and averaged subjective difficulty ratings respectively as dependent variables.

The statistical analyses for reconstruction accuracies and behavioural data were performed using *pingouin*, a statistics package for Python 3 (Vallat, 2018). In case of a significant interaction or a trend, a simple effect test was performed via the Matlab’s Statistics and Machine Learning Toolbox in order to pinpoint the nature of the interaction. Furthermore, comparisons of spectral fine details between face masks and no masks, were computed in Matlab with the *Measures of Effect Size* toolbox (Hentschke & Stüttgen, 2011, see Table S2).

## Results

### Behavioural results

Comprehension performance scores were generated using two ‘true or false’ comprehension questions at the end of each of the 40 trials. We used a two-way repeated measures ANOVA to investigate the influence of the factors *Mask* and *Distractor* on the comprehension performance. Apart from the effect for the distractor speaker (*F(1,28) = 26*.*15, p <*.*001, η*_*p*_^*2*^ *=* .*48)* the results showed no significant influence of face masks (*F(1,28) = 1*.*03, p =* .*32, η*_*p*_^*2*^ *=* .*04*) and no significant interaction (*F(1,28) =* .*02, p =* .*88, η*_*p*_^*2*^ *=* .*001*) between the two factors.

Furthermore, we analysed the subjectively reported difficulty for each condition. We again used two-way repeated measures ANOVA, which showed a significant effect for the distractor speaker (*F(1,28) = 101*.*83, p <* .*001, η*_*p*_^*2*^ *=* .*78*) as well as the face mask (*F(1,28) = 13*.*78, p =* .*001, η*_*p*_^*2*^ *=* .*33*), while not showing a significant effect for the interaction (*F(1,28) = 1*.*33, p =* .*26, η*_*p*_^*2*^ *=* .*06*). These results suggest that, while face masks do not reduce comprehension performance in our setting, they nonetheless lead to a significant increase in perceived listening difficulty.

### Analysis of stimulus reconstruction

Using a stimulus reconstruction approach based on the recorded MEG data, we studied which speech-related features are impaired through face masks, with a special focus on difficult listening situations. We therefore analysed the correlation coefficients (Pearson’s r) obtained using a backward model (Crosse, Di Liberto, Bednar, et al., 2016). The correlation coefficient represents how well the specific stimulus characteristic was reconstructed from brain data and serves as a proxy for how well these features are represented in the neural signal.

With this approach, we generated one correlation coefficient for each condition per participant. This process was repeated for each speech feature of interest. To analyse the effect of the face mask and the distractor speaker, we performed a two-way repeated measures ANOVA, with the Fisher z-transformed correlation coefficients as dependent variables. Detailed results and statistical values are found in the supplementary material (see Table S1). As expected, results show a strong effect (all *p <* .*001*) of the distractor speaker on the stimulus reconstruction across all stimulus characteristics of interest. Figure 2A shows example reconstructions for the speech envelope and the averaged second and third formant (Formant 2/3 or F2/3) as well as mean reconstruction accuracies for clear audiovisual speech (i.e. stimulation material with no mask and no distractor) in Figure 2B.

**Figure 2.**
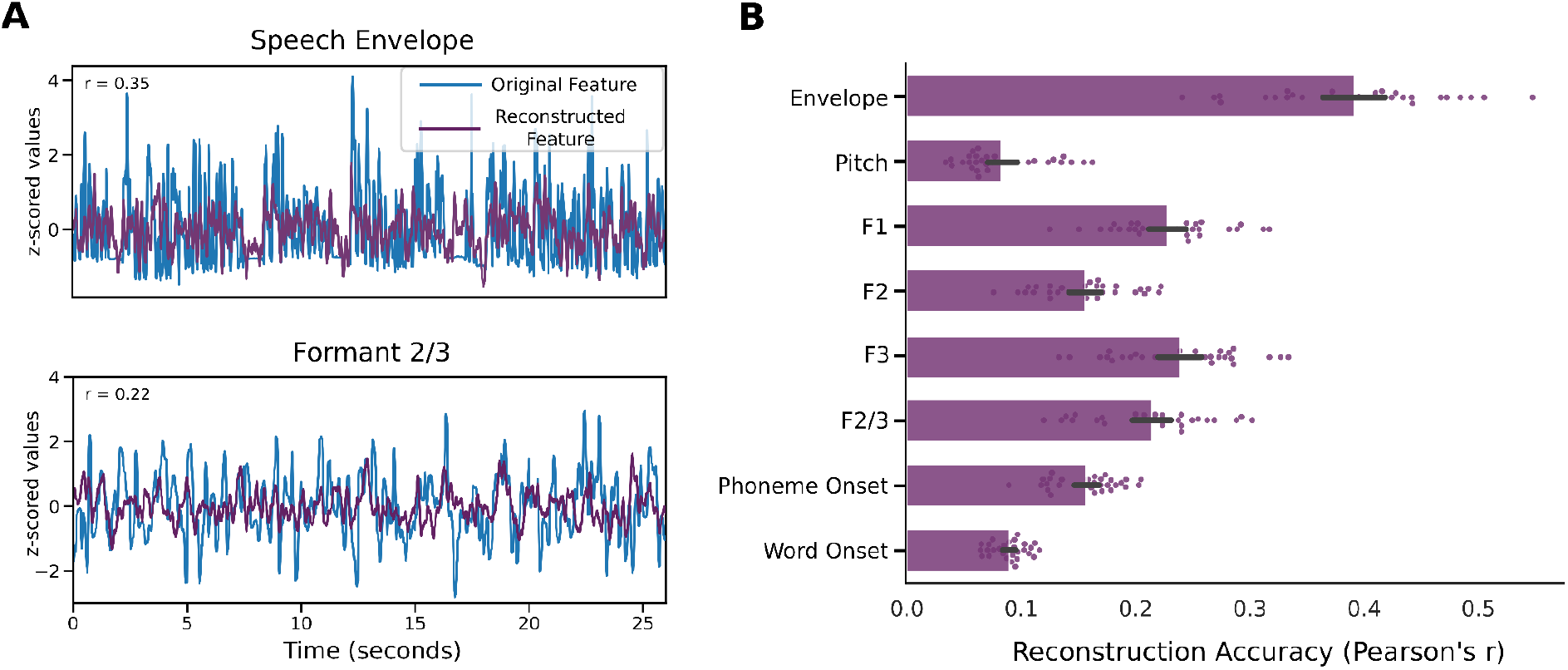
Descriptive depiction of stimulus reconstruction accuracies. **A** Two example stimulus reconstructions of the speech envelope and the averaged F2 and F3 (Formant 2/3, F2/3) for one participant, stimulated with clear audiovisual speech (i.e. stimuli with no mask and no distractor). **B** Mean stimulus reconstruction accuracy for clear audiovisual speech (i.e. stimuli with no mask and no distractor) across participants. Error Bars denote 95% confidence interval.

### Reconstruction of the speech envelope is generally affected by face masks

We investigated how the stimulus reconstruction of the speech envelope is impaired through face masks, with a particular focus on difficult listening situations induced by a distractor speaker. Apart from the negative impact of the distractor speaker (*F(1,28) = 161*.*09, p <* .*001, η*_*p*_^*2*^ *=* .*85*), we observed a strong negative effect of face masks on reconstruction accuracies of the speech envelope (F(1,28) = 24.42, *p* < .001, *η*_*p*_^*2*^ = .47, Figure 3A). We found no significant interaction between the factors *Mask* and *Distractor* (*F(1,28) =* .*25, p =* .*619, η*_*p*_^*2*^ *=* .*01*, Figure 3B and Figure 3C). As the speech envelope conveys crucial information about the syntactic structure of speech (Giraud & Poeppel, 2012; Poeppel & Assaneo, 2020), reduced reconstruction accuracy points to difficulties in deriving this information for further higher-level speech processing.

**Figure 3.**
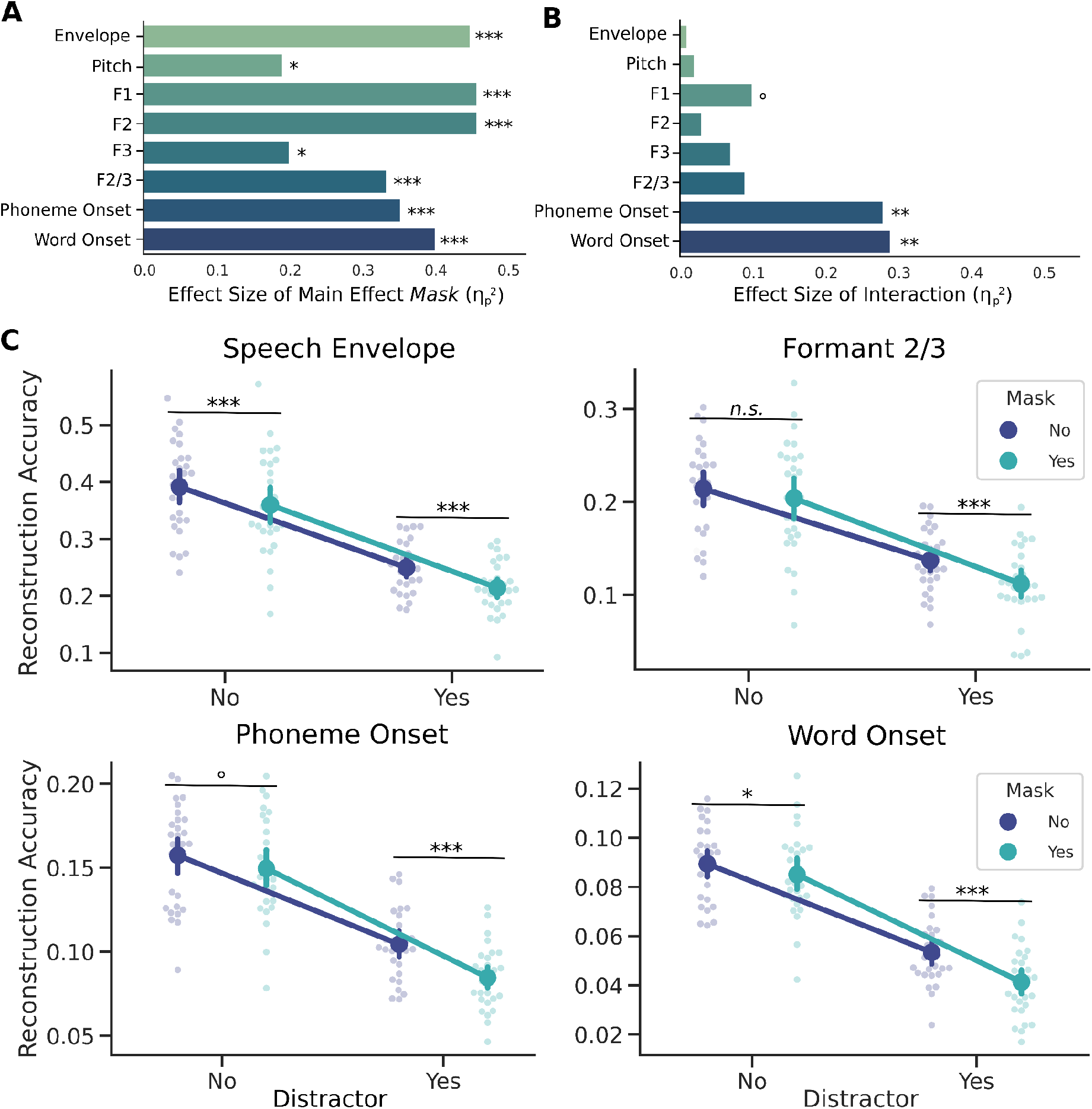
Depiction of the effects of face masks on several speech characteristics. **A** Graphical depiction of the effect size for the main effect of factor *Mask*. Asterisks denote the significance of the effect of the face mask regarding each characteristic. **B** Graphical depiction of the effect size of the interaction of the factors *Mask* and *Distractor*. Asterisks denote the significance of interaction. **C** Depiction of the effects for the speech features speech envelope, averaged F2 and F3 (Formant 2/3, F2/3), and phoneme and word onsets split up for the effects of the face mask and the distractor. Error bars show 95% CI. Asterisks denote the significance of simple effect comparison tests. *n*.*s*.: *p* > .1, ° : *p* < .1, * : *p* < .05, ** : *p* < .01, *** : *p* < .001

### Reconstruction of important spectral fine details is generally affected by face masks

Moreover, we wanted to investigate the influence of face masks on spectral fine details of speech. In this study, we specifically analysed pitch (or fundamental frequency, F0), the first formant (F1), the second formant (F2) and the third formant (F3). When facing concurrent speakers, a listener must segregate the speech signal into different speech streams. It is suggested that pitch serves a fundamental role in this process (Bregman, 1990). Formants are especially interesting, as they are vital for identifying vowels (Peterson & Barney, 1952) which are the voiced components of speech. Additionally, we investigated the averaged F2 and F3 (F2/3), as these two formants generated in the front cavity converge into ‘focal points’ after specific vowel-consonant combinations (Badin et al., 1990) and their frequency range has been shown to correlate strongly with lip movements (Chandrasekaran et al., 2009). Furthermore, this speech feature has been shown to be tracked by visual-only speech and is therefore prone to be affected through face masks (Suess, Hauswald, et al., 2021). Detailed results for F1, F2, F3 are depicted in Table S1 (see Supplementary Material). Effect sizes of the main effect are presented graphically in Figure 3A and for the interactions in Figure 3B. With a distractor speaker, reconstruction of pitch (*F(1,28) = 89*.*18, p <* .*001, η*_*p*_^*2*^ *=* .*76*) and F2/3 (*F(1,28) = 75*.*81, p <* .*001, η*_*p*_^*2*^ *=* .*73*) was reduced. The reconstruction of the pitch shows an impairment through face masks (*F(1,28) = 7*.*26, p =* .*018, η*_*p*_^*2*^ *=* .*21*) with no significant interaction (*F(2,28) =* .*49, p =* .*487, η*_*p*_^*2*^ *=* .*02*). The same is true for F2/3, which showed a significant reduction of reconstruction accuracy through face masks (*F(1,28) = 14*.*78, p <* .*001, η*_*p*_^*2*^ *=* .*35*). While reconstruction of F2/3 was not affected through a face mask when no distractor was present (No Distractor: *MD (SE) =* .*011 (*.*006), p =* .*107*), it was reduced in presence of a distractor (Distractor: *MD (SE) =* .*025 (*.*007), p <* .*001*). This interaction was however not significant (*F(1,28) = 2*.*76, p =* .*108, η*_*p*_^*2*^ *=* .*09*). These results suggest that face masks do impair the tracking of spectral fine details of the speech relevant spectrum generally irrespective of a distractor speaker.

### Reconstruction for phonetic and lexical boundaries is impaired through face masks specifically in difficult listening situations

Detecting lexical boundaries is important for chunking the continuous speech stream into meaningful interpretable units. As a last step, we therefore investigated how face masks impair the reconstruction of phoneme and word onsets.

For phoneme onsets, we found significant main effects of reconstruction accuracies for the factor *Distractor (F(1,28) = 187*.*81, p <* .*001, η*_*p*_^*2*^ *=* .*87*) and *Mask (F(1,28) = 16*.*63, p <* .*001, η*_*p*_^*2*^ *=* .*37*), as well as a strong significant interaction of *Mask* and *Distractor* (*F(1,28) = 10*.*75, p =* .*003, η*_*p*_^*2*^ *=* .*28*). Similar results can be shown for word onset reconstruction accuracies with significant main effects of the *Distractor* (*F(1,28) = 278*.*19, p <* .*001, η*_*p*_^*2*^ *=* .*91*), *Mask* (*F(1,28) = 19*.*95, p <* .*001, η*_*p*_^*2*^ *=* .*42*) and the interaction (*F(1,28) = 11*.*46, p =* .*002, η*_*p*_^*2*^ *=* .*29*). For phoneme onset, post-hoc simple effect tests revealed significant differences for the factor *Mask* when a distractor was present, while only showing a trend when no distractor was presented (No Distractor: *MD (SE) =* .*008 (*.*004), p =* .*058*; Distractor: *MD (SE) =* .*020 (*.*004), p <* .*001*, see Figure 3C). For word onsets, we found significant differences irrespective of a distractor speaker, but a strongly increasing effect when a distractor speaker was presented alongside (No Distractor: *MD (SE) =* .*004 (*.*002), p =* .*017*; Distractor: *MD (SE) =* .*012 (*.*003), p <* .*001*). Following this, face masks seem to decrease the ability to segment the speech stream into meaningful units when listeners are encountering challenging listening situations.

## Discussion

The effects of face masks on speech comprehension have been investigated in various studies on a behavioural level (Brown et al., 2021; Giovanelli et al., 2021; Rahne et al., 2021; Toscano & Toscano, 2021; Yi et al., 2021). Despite the overall agreement of adverse effects of face masks on speech comprehension, it has been unclear which features of speech processing are specifically affected.

Our results show that tracking of features responsible for successful processing of naturalistic speech is impaired through (surgical) face masks. From general temporal modulations of the speech envelope to modulations of spectral fine details (pitch and formants) and segmentation of speech (phoneme and word onsets), a face mask significantly reduces the decodability of these features from brain data. However, not all of these speech features are affected by the face mask the same way. While the brain’s tracking of low-level acoustic features (i.e. the speech envelope and spectral fine details) are affected generally, the higher-level segmentational features phoneme onset and word onset show particularly strong reduction of reconstruction accuracy through face masks when facing a challenging listening situation (i.e. using a distractor speaker).

### Occlusion of lip movements increases subjective listening difficulty, while speech comprehension is unaffected

Regarding our behavioural results, we observed significantly decreased performance through a distractor speaker, but not through the face mask. This is in line with previous findings on audio-only speech (Toscano & Toscano, 2021) which found no significant effect of surgical face masks on word recognition in easy and challenging listening situations. However, another study with audiovisual speech found significant effects of a surgical face mask in conditions of moderate (−5 dB SNR) and high (−9 dB SNR) background pink noise on sentence intelligibility (Brown et al., 2021). As our study used longer duration audiobooks, our behavioural measurements might not have been precise enough (i.e only two binary unstandardised ‘true or false’ statements at the end of each trial regarding semantic comprehension) to detect this influence.

We also found that subjective ratings of listening difficulty were significantly larger when speakers wore a face mask independent of a distractor speaker. An explanation for this is that removing informative visual cues leads to an increase of linguistic ambiguity resulting in more effortful mental correction by the listener (Hughes et al., 2018). This increased effort however might be at the same time compensating the influence of face masks on the aforementioned comprehension performance (Winn & Teece, 2021). Despite a comparable performance in speech comprehension between conditions with- and without masks, listening to a speaker wearing a mask increases the subjective listening effort. Such increased effort has been associated with social withdrawal in the hearing impaired population (Hughes et al., 2018) and should not be disregarded. Still, our behavioural results contradict previous findings, which only showed an effect of face masks on listening effort when combined with background noise (Brown et al., 2021). Again, differences in study design (one minute audiobooks vs. single sentence) may account for this difference.

### Occlusion of lip movements impairs tracking of crucial acoustic speech features generally

The speech envelope, mostly associated with conducting syntactic and phonetic information (Giraud & Poeppel, 2012; Poeppel & Assaneo, 2020), has been deemed a core characteristic regarding speech tracking (Brodbeck & Simon, 2020). In multi-speaker listening situations, attending to the target speaker is related to enhanced tracking of the envelope of the attended speech compared to the unattended speech (O’Sullivan et al., 2015; Park et al., 2016; Zion Golumbic et al., 2013). A reduced tracking of this speech feature might represent a difficulty in following and segmenting the target speech stream when confronted with face masks. Also using AV speech, a study showed the audiovisual benefits for speech envelope tracking, especially in the context of background noise (−9 SNR) (Crosse, Di Liberto, & Lalor, 2016). Our results do not confirm this effect of inverse effectiveness for the tracking of the speech envelope and other acoustic features. Instead, we see a strong effect hinting at visual effects independent of acoustic noise. These differences might be explained by differences in study design, as we used a distractor speaker at 0 dB SNR compared to background noise at -9 dB SNR. While not confirming the notion of inverse effectiveness, our results are in line with a study by (Golumbic et al., 2013) showing an increased response in the auditory cortex to AV stimuli compared to audio-only stimuli irrespective of a distractor speaker. Their results point at a modulatory influence of the visual speech on the auditory processing.

Regardless of the importance of the speech envelope, it does not convey specific information regarding certain phonetic objects, like vowels and vowel-consonant combinations. Formants on the other side define vowels directly (Peterson & Barney, 1952).

While the first (F1) and (F2) are generally considered core formants in speech (Peterson & Barney, 1952), using an averaged F2 and F3 (F2/3) instead of F2 has proven to be beneficial as it smooths transitions from one vowel to the other (Stevens, 2000) and due to their convergence in the front cavity (Badin et al., 1990). With regards to visual speech tracking, the encompassed frequencies of F2 and F3 correlate strongly with lip movements (Chandrasekaran et al., 2009) so that these frequencies likely contribute to a visual-phonological transformation (cf. Hauswald et al. (2018)). While Hauswald et al. (2018) proposed a role of the visually conveyed envelope information for a visuo-phonological transformation, another study by our group further suggests that also the visually transported formant information go through such a transformation (Suess, Hauswald, et al., 2021), which is possibly even more relevant for visual speech processing than the transformation of the speech envelope (Plass et al., 2020). Finally, the reconstruction of voice pitch or fundamental frequency, used to segregate concurrent speech streams (Bregman, 1990), is also reduced through face masks, which might lead to difficulties disentangling the target speech stream and the distractor speech stream. Taking the effects face masks have on the envelope, pitch and formants together, face coverings might lead to subsequent difficulties in identifying phonemes and as a consequence also words. As we used surgical face masks in our study, which have a small influence on the speech acoustics and attenuate only higher frequencies above 3 kHz (Corey et al., 2020; Toscano & Toscano, 2021) and found only small differences between stimuli with and without face mask (see Table S2), we attribute these findings mostly to the missing visual input. This is further supported by the fact that investigated spectral fine details (namely pitch and formants) present themselves in frequencies below 3 kHz (Peterson & Barney, 1952). We therefore interpret the aforementioned effects as the result of a missing visual input and a subsequent impossibility to integrate acoustic and visual information in contrast to effects of distorted acoustics through the surgical face mask.

### Occlusion of lip movements impairs tracking of higher-level segmentational features especially in challenging listening situations

Tracking of phoneme and word onsets is affected such that face masks impair chunking in challenging listening situations especially strong. Studies investigating simple ERPs when listening to continuous speech found enhanced responses to word onsets (Sanders et al., 2002; Sanders & Neville, 2003), pointing to an internal chunking mechanism of the brain for optimal speech processing. On a lower level, brain responses induced by phoneme onsets are reliably predicted by encoding models (Brodbeck et al., 2018; Daube et al., 2019; Di Liberto et al., 2015), implying chunking already on this level. When deprived of visual cues (through face masks) and in noisy acoustic environments, our findings suggest that individuals face problems with segmenting the continuous speech stream into meaningful units (i.e. words and phonemes). Furthermore, formant frequencies might also play an important role in detecting syllables and more importantly phonemes and their boundaries (Plass et al., 2020). For compensating this degradation in challenging listening situations, watching the speaker’s face provides important information (Mitchel & Weiss, 2014) for word segmentation. Highlighting this even further, visual cues from the mouth area have been found to enhance phonetic discrimination, by providing visemic information (Fisher, 1968). Taken together, depriving listeners of these visual cues through covering the mouth affects an important step of unit identification (words and phonemes), which helps chunking the stream for further processing. These results confirm that multisensory gain is increased when individual input from a modality is weak or distorted (Crosse, Di Liberto, & Lalor, 2016). Interestingly, in our results we only find this effect for higher-level segmentation features, while all lower level features show general effects of the face mask.

With this study, we expand the knowledge about multisensory AV speech processing by putting past findings into the context of face masks. Expectations about the influence of face masks on speech characteristics were confirmed in the way that it impairs stimulus feature reconstruction in difficult listening situations, following the concept of inverse effectiveness. This effect can be shown in higher-level features of speech segmentation (i.e. phoneme onsets and word onsets) in the form of an interaction between the face mask and the distractor speaker, while reconstruction of acoustic information is generally impaired. This could again point to a visuo-phonological transformation process from the visual input to a phonetic representation in the range of F2 and F3, which is however not possible when speakers wear a face mask. The phonetic representation in the visual cortex might be influencing further processing of the complex speech signal in the auditory cortex through direct pathways from visual to auditory regions (Besle et al., 2004, 2009; Golumbic et al., 2013) or by influencing connectivity between the auditory cortex and higher-level regions specialized for speech processing (Giordano et al., 2017). Another possible inhibited process through the visual occlusion of the mouth area was presented in a recent study. They provided evidence for a linguistic representation in the visual cortex stemming from visemic information of speech, a process independent of auditory processing associated with lip reading (Nidiffer et al., 2021).

### Practical implications

Following our findings, the use of transparent face masks is principally favourable. However, some of the current transparent models come with significantly reduced transmission of acoustic detail (Corey et al., 2020) resulting in reduced intelligibility and increased difficulty ratings, when presented in noisy environments (Brown et al., 2021). It is also important to consider that this study investigated normal hearing subjects, and results for individuals with hearing loss might be different (Puschmann et al., 2019). In line with this notion, data collected before the Covid-19 pandemic suggests strong benefits of transparent face masks for listeners with hearing loss (Atcherson et al., 2017) allowing them to integrate visual information for speech processing. A recent study confirms this by comparing the impact of surgical face masks to (transparent) face shields (Homans & Vroegop, 2021). Despite the face shield’s larger impact on acoustics compared to the surgical face masks, individuals with hearing loss showed no significant decrease in speech intelligibility when confronted with a face shield compared to no facial covering, while scores were significantly worse when a surgical face mask was worn.

## Conclusion

With this study, we investigated the effects of face masks on multisensory processing of speech. Using a stimulus reconstruction approach, we found a general impairment of AV speech integration through face masks, while higher-level segmentational feature effects were most pronounced when a distractor alongside a face mask wearing target speaker was presented. The latter finding is in line with the concept of inverse effectiveness, suggesting increased multisensory gain in the context of a weak single modality, which is however not possible when the lip movements are occluded. Here, we can therefore show differential effects for low level acoustics (envelope, pitch and formants) and higher level segmentational features of speech. Our results strengthen the concept of a visual-phonological transformation improving tracking of speech features through visual information. The present findings might have important implications for deciding which face masks to use, especially when dealing with the hearing impaired.

## Acknowledgements

This work is supported by the Austrian Science Fund, P31230 (“Audiovisual speech entrainment in deafness”) and P34237 (“Impact of face masks on speech comprehension”). Sound icon made by *Smashicon* from www.flaticon.com.

Thanks to the whole research team. Special thanks to Fabian Schmidt for providing support for graphical design.

## Code Availability

The code to recreate figures and statistics is available at the corresponding author’s GitLab repository (https://gitlab.com/CLH96/facemasktracking). Further and more detailed information regarding the method can be provided upon request.

## Supplementary Material

**Table S1.** Results of main effects and interaction for reconstruction accuracy.

|  | Main effect: *Mask* |  |  | Main effect: *Distractor* |  |  | Interaction |  |  |
| --- | --- | --- | --- | --- | --- | --- | --- | --- | --- |
| | F(1,28) | *p* | $\eta_p^2$ | F(1,28) | *p* | $\eta_p^2$ | F(1,28) | *p* | $\eta_p^2$ |
| Envelope | **24.42** | **<.001** | **.47** | **161.09** | **<.001** | **.85** | .25 | .619 | .01 |
| Pitch | **7.26** | **.018** | **.21** | **89.18** | **<.001** | **.76** | .49 | .487 | .02 |
| F1 | **25.70** | **<.001** | **.48** | **146.68** | **<.001** | **.84** | 3.15 | .087 | .10 |
| F2 | **25.38** | **<.001** | **.48** | **50.13** | **<.001** | **.64** | .77 | .388 | .03 |
| F3 | **7.36** | **.013** | **.21** | **99.90** | **<.001** | **.78** | 2.18 | .152 | .07 |
| F2/3 | **14.78** | **<.001** | **.35** | **75.81** | **<.001** | **.73** | 2.76 | .108 | .09 |
| Phoneme Onset | **16.63** | **<.001** | **.37** | **187.81** | **<.001** | **.87** | **10.75** | **.003** | **.28** |
| Word Onset | **19.95** | **<.001** | **.42** | **278.19** | **<.001** | **.91** | **11.46** | **.002** | **.29** |

Note. Bold letters indicate significant effects.

**Table S2.** Mean frequencies of spectral fine details.

|  | Female |  | Male |  | Statistics |  |  |
| --- | --- | --- | --- | --- | --- | --- | --- |
|  | *No mask* | *Mask* | *No Mask* | *Mask* | t(39) | *p* | *g* |
| Pitch | 205.8 (6.1) | 203.0 (9.2) | 119.9 (6.6) | 120.8 (5.9) | .79 | .435 | .02 |
| F1 | 687.1 (14.2) | 690.4 (16.8) | 663.0 (20.1) | 631.1 (16.4) | 1.89 | .065 | .14 |
| F2 | **1906.1 (22.3)** | **1878.3 (23.2)** | **1697.5 (26.8)** | **1668.8 (16.6)** | **11.89** | **<.001** | **.26** |
| F3 | **2954.1 (14.1)** | **2911.6 (19.9)** | **2604.7 (27.1)** | **2551.9 (19.6)** | **14.01** | **<.001** | **.26** |
| F2/3 | **2427.3 (16.9)** | **2392.1 (20.4)** | **2149.1 (25.7)** | **2108.1 (16.4)** | **14.00** | **<.001** | **.26** |

Note. Numbers in parentheses denote the standard deviation. Bold letters indicate significant differences. For descriptive purposes, the mean frequencies are split up for male and female speakers. The statistical analysis was performed using paired sample t-test comparing features (pitch and formants) extracted out of the 40 stimuli with no mask to their counterpart with speakers wearing a mask.

